# Natural killer-like B cells promote Th17 cell differentiation and exacerbate rheumatoid arthritis

**DOI:** 10.1101/2021.10.27.466096

**Authors:** Ping Wang, Jing Song, Mingxin Bai, Xi Zheng, Yang Xie, Yundi Tang, Xin Li, Xiangyu Fang, Yuan Jia, Limin Ren, Hua Ye, Yin Su, Shuo Wang, Zhanguo Li, Fanlei Hu

## Abstract

B cells are important participants in the pathogenesis of rheumatoid arthritis (RA). Besides classical B cells, novel B cell subsets are continually to be identified in recent years. Natural killer-like B (NKB) cells, a newly recognized B cell subset, are proved to be actively involved in the anti-infection immunity. However, their role in RA and the potential mechanism remain elusive. Here, we showed that NKB cells were expanded dramatically in collagen-induced arthritis (CIA) mice, demonstrating dynamic changes during the disease progression. These cells promoted CD4^+^ effector T cell proliferation and Th17 cell differentiation *in vitro*, while adoptive transfer of these cells exacerbated the arthritis severity of CIA mice. RNA Sequencing revealed that NKB cells displayed distinct differential gene expression profile under RA circumstance, potential perpetuating the disease progression. Moreover, the frequencies of NKB cells were significantly increased in RA patients, positively correlated with the clinical and immunological features. After effective therapy, these cells could be recovered to normal levels. Taken together, our results preliminarily revealed the pathogenic role of NKB cells in RA by promoting Th17 proinflammatory responses. Targeting these cells might provide potential therapeutic strategies for this persistent disease.

## Introduction

Rheumatoid arthritis (RA) is a common chronic inflammatory autoimmune disease characterized by synovitis, pannus formation, cartilage, and bone destruction, with various immune cells involved, including T cells, B cells, NK cells, DC cells, myeloid-derived suppressor cells, and so on(Guo et al., 2016; Smolen, Aletaha, & McInnes, 2016). Although the pathogenesis of RA remains to be fully elucidated, B cells are proved to be the major participants in the initiation and perpetuation of RA(Bugatti, Vitolo, Caporali, Montecucco, & Manzo, 2014).

B cells play crucial roles both in innate and adaptive parts of the immune system. Despite the maturity of the field of B cell biology, there are still many completely unexpected things to be discovered(Tarlinton, 2019). Moreover, B cells play an important role in the pathogenesis of autoimmune diseases. The pathogenic roles of B cells involved in the adaptive immunity of RA have been studied in-depth, including producing autoantibodies, presenting auto-antigens, and secreting cytokines(Marston, Palanichamy, & Anolik, 2010). The impairment of regulatory B cells (Bregs) has also been recognized in the development of RA, including IL-10-producing Breg (B10), GrB-producing Breg, and so forth(Daien et al., 2014; F. Hu et al., 2017; Ummarino, 2017; Xu et al., 2017). In addition, the innate-like B cells (ILBs), whose cell-surface markers, tissue distribution, and functions are different from those of conventional B cells, blur the boundaries of innate immunity and adaptive immunity and are also actively involved in RA pathogenesis(Cerny & Striz, 2019; F. L. Hu et al., 2018; Romero-Ramírez et al., 2019). More and more studies revealed that B cells are a heterogenous population with distinct functions. Further identifying novel B cell subsets will help better understanding their role in the pathogenesis of RA, and benefit the targeted therapy.

ILBs are heterogeneous B cell populations with innate sensing and responding properties(Grasseau et al., 2020; Zhang, 2013). Different ILBs share similarities such as rapid response potential, restricted BCR repertoires, and regulatory roles in innate and adaptive immunity(Zhang, 2013). In addition to B-1a cells and marginal zone B cells, innate response activator B cells, T-bet positive B cells, IL-17-producing B cells, human self-reactive VH4-34-expressing B cells, and natural killer-like B (NKB) cells are all termed as ILBs(Tsay & Zouali, 2018).

NKB cells are a newly identified ILB population both in mice and human(Wang et al., 2016). Mouse NKB cells express CD19^+^NK1.1^+^ while human express CD19^+^NKp46^+^ surface markers. They exhibit unique signature features that are distinct from those of NK cells and conventional B cells(Wang, Xia, & Fan, 2017). NKB cells show a great potential to produce IL-18 and IL-12 at the early stage of infection, leading to activation of type 1 innate lymphoid cells (ILC1s) and NK cells against microbial infection(Wang et al., 2016). Studies showed that the number of NKB cells in the spleen and the level of IL-18 in serum were markedly increased in experimental alcoholic liver injury, which could be reversed after treatment(Ge et al., 2019). NKB cells were also found to systemically distribute in rhesus macaques, and expand in the peripheral blood and colon during SIV infection(Manickam et al., 2018). However, the characteristics and their potential role in RA remain unclear.

In this study, by studying both collagen-induced arthritis (CIA) mice and RA patients, we investigated the involvement of NKB cells and the underlying mechanisms in RA development.

## Results

### NKB cells are expanded in autoimmune arthritis mice and show dynamic changes during the disease progression

We first detected the distribution of NKB cells in mouse tissues. The results showed that NKB cells were widely distributed in DBA/1 mice, including spleen, mesenteric lymph node (MLN), draining lymph node (DLN, inguinal lymph node, popliteal lymph node, axillary lymph node, and submandibular lymph node), Peyer patch, and paw, with spleen as the dominant enriching site (Figure 1A-C).

**Figure 1.**
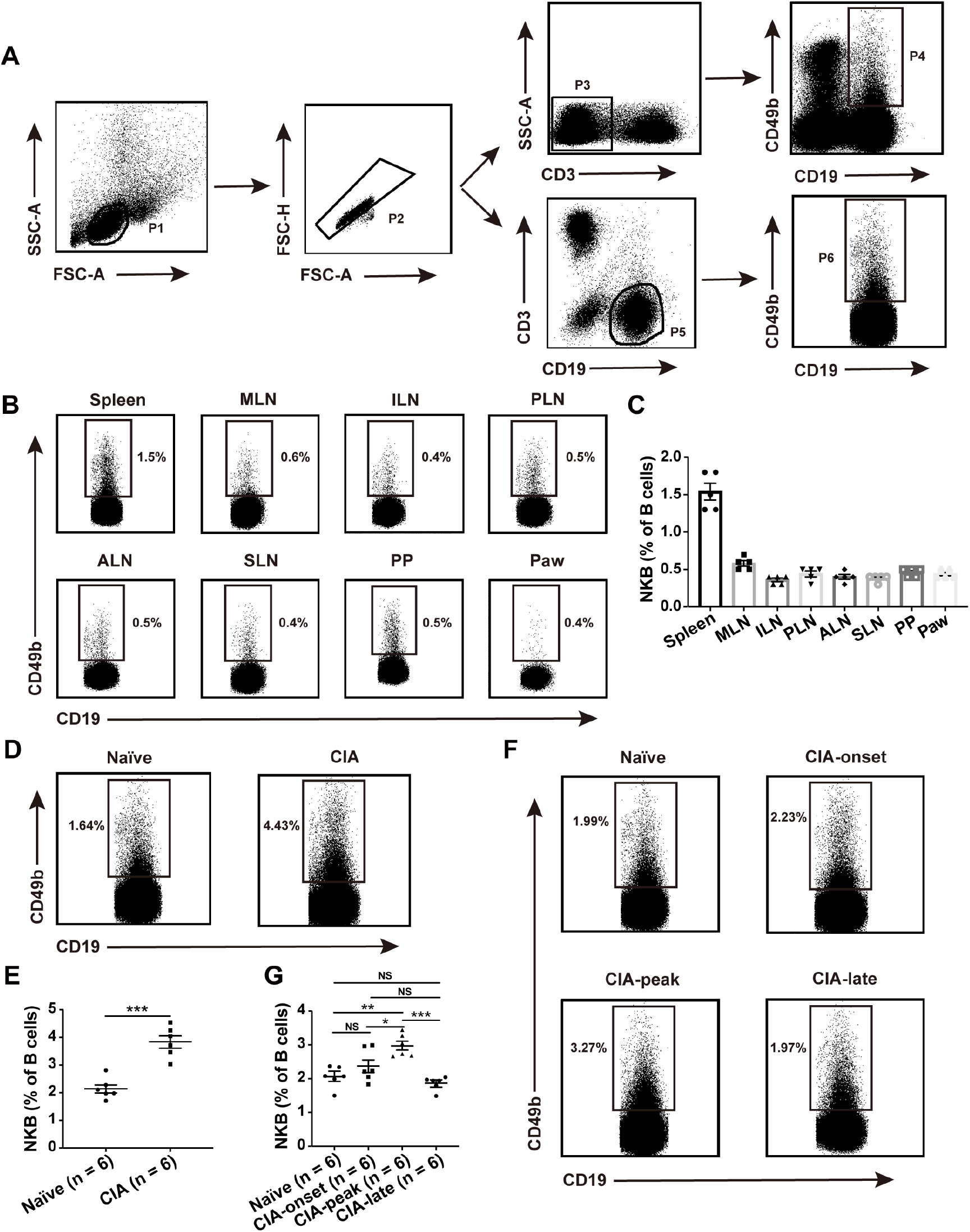
Distribution of NKB cells in mice and their dynamic changes during the development of collagen-induced arthritis (CIA). (**A**) Gating strategy for mouse NKB cells. (**B, C**) Distribution of NKB cells in DBA/1 mouse tissues, including spleen, mesenteric lymph node (MLN), inguinal lymph node (ILN), popliteal lymph node (PLN), axillary lymph node (ALN), submandibular lymph node (SLN), Peyer patch (PP), and paw. The representative flow charts (**B**) and the statistical result (**C**) were shown, respectively. (**D, E**) Flow cytometric analysis of the frequencies of NKB cells in the spleen of naïve and CIA mice (n = 6 per group). The representative flow charts (**D**) and the statistical result (**E**) were shown, respectively (two-tailed student’s *t*-test, \*\*\**P* < 0.001). (**F, G**) Flow cytometric analysis of the frequencies of NKB cells during the development of CIA. Spleens from naïve, CIA-onset (< 2 days), CIA-peak (∼ 15 days), and CIA-late (> 60 days) mice were collected for NKB detection (n = 6 per group). The representative flow charts (**F**) and the statistical result (**G**) were shown (one-way ANOVA test followed by Tukey’s posttest for multiple comparisons, \**P* < 0.05, \*\**P* < 0.01, \*\*\**P* < 0.001, NS, not significant). Data are presented as mean ± SEM. Results are representative of three independent experiments.

We then constructed the commonly used arthritis model, CIA in DBA/1 mice, and detected the frequencies of NKB cells during the disease development. It was shown that NKB cells were significantly expanded in the spleen of CIA mice as compared with naïve mice (Figure 1D, E). To further characterize the dynamic changes of NKB cells during arthritis progression, we detected their frequencies in different stage CIA mice. According to the time post arthritis initiation, CIA mice were divided into the following three stages: onset stage (< 2 days), peak stage (∼15 days), and late-stage (> 60 days). It was shown that the frequencies of NKB cells were slightly increased at the onset stage, and were significantly upregulated at the peak stage, then were recovered to normal levels at the late stage (Figure 1F, G). Moreover, NKB precursor (NKBP) cells in the bone marrow, defined as Lin^-^CD122^+^ CD19^+^ CD49b^+^ cells, were also detected during the CIA development. Interestingly, it was found that the frequencies of NKBP cells at the onset and late stage of CIA mice were significantly decreased (Supplementary Figure 1A, B), probably due to their enhanced maturation and transportation to the spleen. Nevertheless, the detailed mechanisms need to be further studied.

### NKB cells promote CD4^+^ effector T cell proliferation and Th17 cell differentiation

We next tried to reveal the role of NKB cells in the pathogenesis of RA. NKB cells and CD4^+^CD25^-^ effector T cells were isolated from the spleens of CIA mice, the purities of which detected by flow cytometry were over 90% (Figure 2A, Supplementary Figure 2A, B), and then subjected to the co-culture assay *in vitro*. The results showed that after co-culture with NKB cells, the proliferation of effector T cells was significantly increased as compared with the control group (Figure 2B-D). Moreover, the frequencies of Th17 cells were increased remarkably in the co-culture group (Figure 2E, F), with the upregulated secretion of IL-17A in the culture supernatants (Figure 2G). Nevertheless, no obvious change was shown for Th1 cells and IFN-γ secretion after co-culture (Figure 2H-J). Furthermore, NKB cells exacerbated the Th17 cell responses in a paracrine and cell-cell contact-dependent manner. Under the transwell co-culture system, NKB cells-mediated enhancement of Th17 cell responses was diminished (Figure 2K-M).

**Figure 2.**
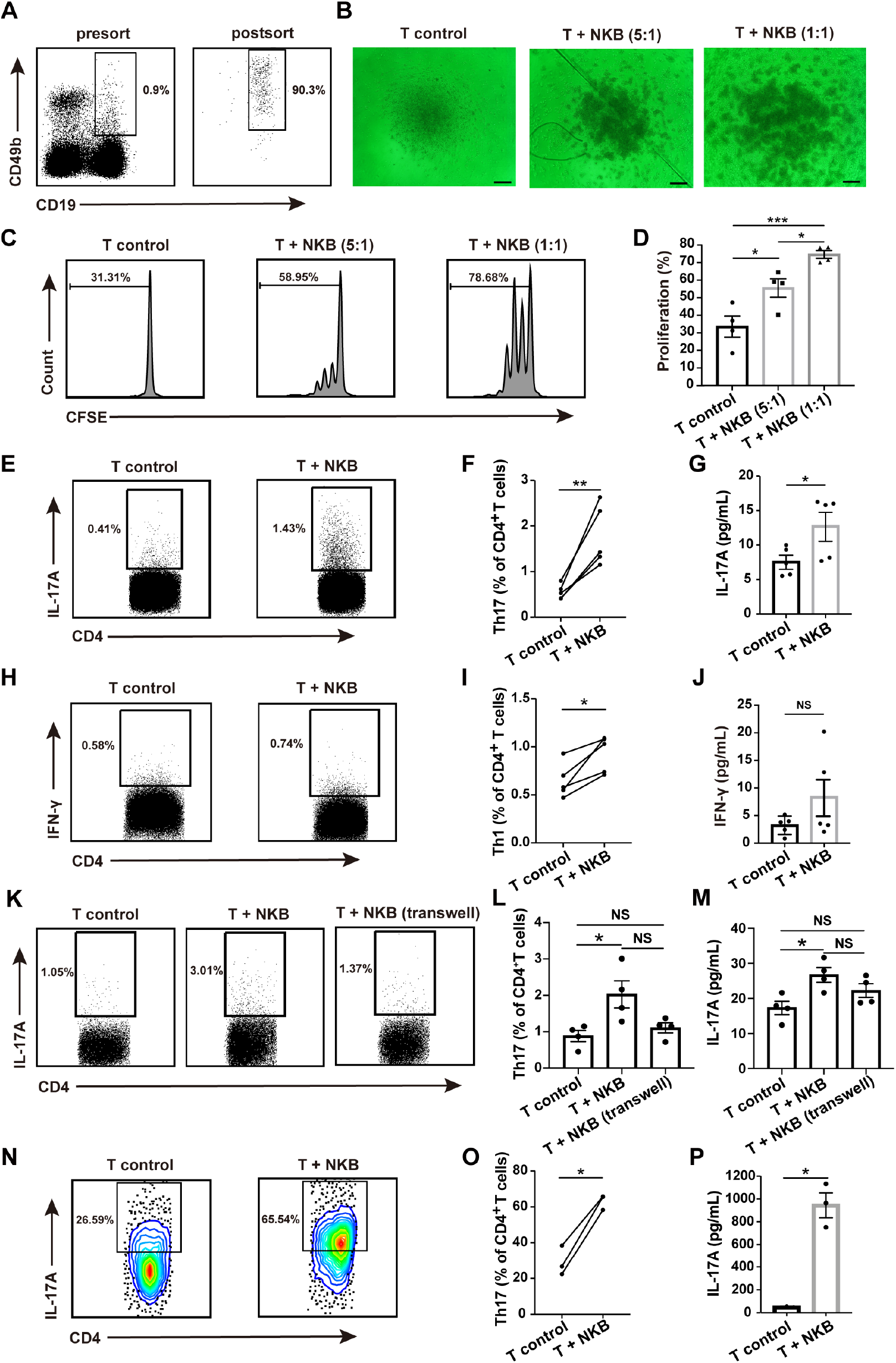
NKB cells promote CD4^+^ effector T cell proliferation and Th17 cell differentiation. CD4^+^CD25^-^ T cells or CD4^+^CD25^-^CD62L^hi^ naïve T cells and NKB cells were isolated from the spleen of CIA mice for co-culture assays. (**A**) The representative flow charts for presort and postsort of mouse NKB cells. (**B-D**) 5×10^4^ CD4^+^CD25^-^ T cells were stained with CFSE and then co-cultured with different numbers of NKB cells (T: NKB = 5:1, or 1:1) *in vitro* for 3 days (n = 4). The representative microscopic images of the cultured cells were shown. Scale bar = 100 μm (**B**). The T cell proliferation was also detected by flow cytometric analysis of the CFSE dilution (**C-D**). The representative flow charts (**C**) and the statistical result (**D**) were shown, respectively (one-way ANOVA test followed by Tukey’s posttest for multiple comparisons, \**P* < 0.05, \*\*\**P* < 0.001, NS, not significant). (**E-J**) 5×10^5^ CD4^+^CD25^-^ T cells were co-cultured with 1×10^5^ NKB cells (5:1) *in vitro* for 3 days (n = 5), then the frequencies of Th17 (**E, F**) and Th1 cells (**H, I**) were detected by flow cytometry, while the levels of IL-17A (**G**) and IFN-γ (**J**) in the cell culture supernatants were detected by ELISA (two-tailed paired *t-*test and two-tailed student’s *t*-test, \**P* < 0.05, \*\**P* < 0.01, NS, not significant). (**K-M**) 5×10^5^ CD4^+^CD25^-^ T cells were co- cultured with 1×10^5^ NKB cells (5:1) under normal or transwell culture systems *in vitro* for 3 days (n = 4), then the frequencies of Th17 cells (**K, L**) were detected by flow cytometry, while the levels of IL-17A in the cell culture supernatants (**M**) were detected by ELISA (one-way ANOVA test followed by Tukey’s posttest for multiple comparisons, \**P* < 0.05, NS, not significant). (**N-P**) 5×10^5^ CD4^+^CD25^-^CD62L^hi^ naïve T cells and 1×10^5^ NKB cells were co-cultured *in vitro* (5:1) for 3 days under the Th17 cell polarization condition (n = 3). The frequencies of Th17 cells (**N, O**) were detected by flow cytometry, while the levels of IL-17A in the cell culture supernatants (**P**) were detected by ELISA (two-tailed paired *t-*test and two-tailed student’s *t*-test, \**P* < 0.05). Data are presented as mean ± SEM. Results are representative of three independent experiments.

To further reveal the effects of NKB cells on Th17 cell differentiation, we co-cultured CD4^+^CD25^-^CD62L^hi^ naïve T cells and NKB cells isolated from the spleens of CIA mice under the Th17 cell polarization conditions. As shown in Figure 2N-P, the frequencies of Th17 cells and the levels of IL-17A in the culture supernatants were both elevated notably in the co-culture group.

Taken together, these results suggest that NKB cells might play a pathogenic role in RA by promoting CD4^+^ effector T cell proliferation and Th17 cell differentiation and responses.

### NKB cells exacerbate autoimmune arthritis in mice

To illustrate the pathogenic role of NKB cells in *vivo*, we adoptively transferred these cells from peak stage CIA mice to the onset stage CIA mice with equal arthritis scores. The scheme of the experimental setup was demonstrated in Figure 3A. The results showed that compared with those mice transferred with saline, the mice transferred with NKB cells demonstrated significantly higher arthritis scores (Figure 3B). NKB cell transfer also led to more severe bone destruction as assessed by micro-CT (Figure 3C) and more serious inflammation as revealed by HE staining (Figure 3D) in the CIA mouse joints. Moreover, NKB cell transfer resulted in the increased frequencies of Th17 cells and Th1 cells in the CIA mouse spleen (Figure 3E-H). However, no obvious changes were shown for Treg cells, Tfh cells, germinal center B cells, and B10 cells after the transfer (Supplementary Figure 3, 4). All these results further confirmed the effects of NKB cells in exacerbating RA.

**Figure 3.**
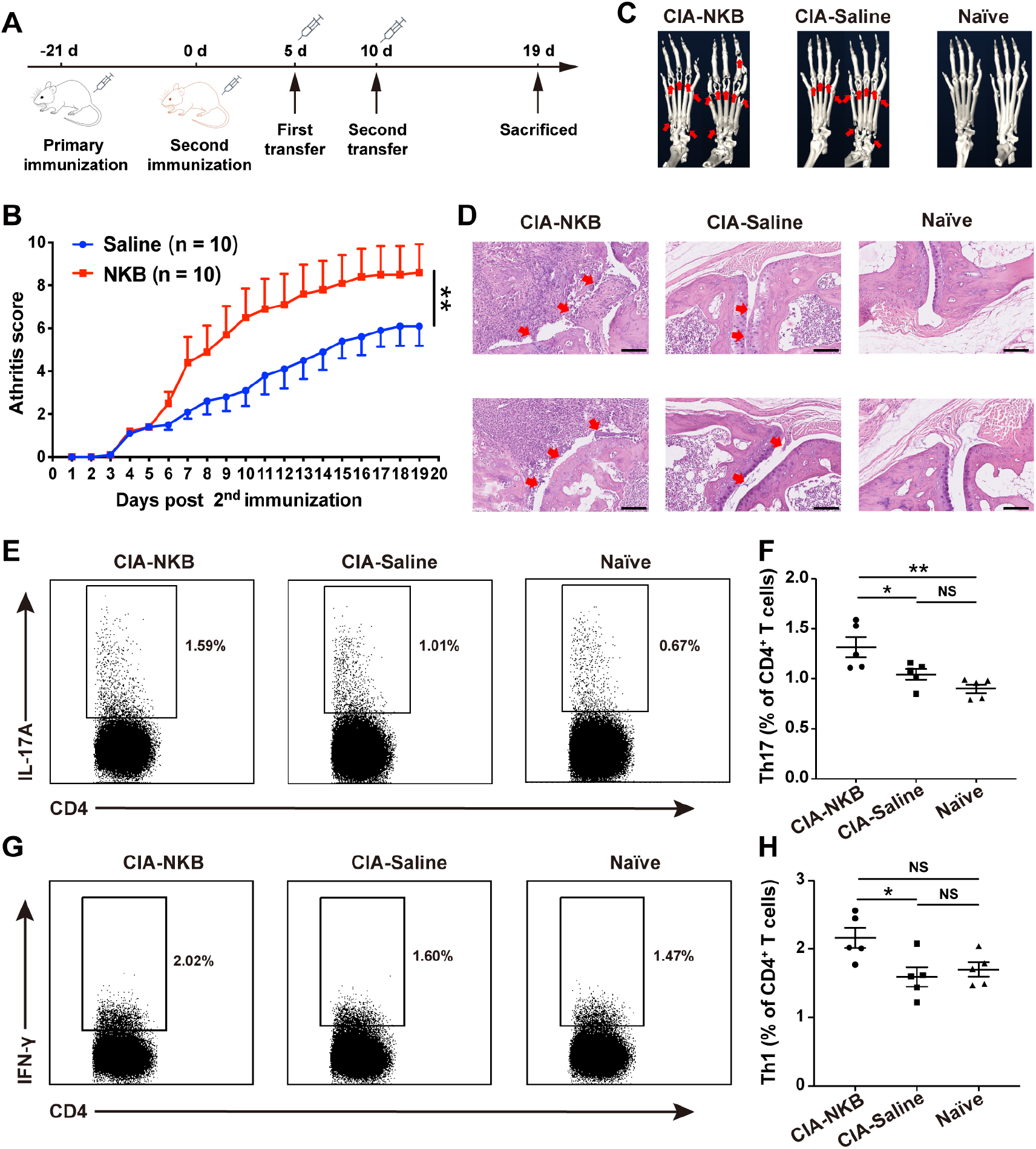
NKB cells exacerbate the arthritis severity in CIA mice. (**A**) Scheme of the experimental setup. CIA mice were established by the first immunization on day - 21 and the second immunization on day 0. Then CIA mice with an average arthritis score of 2 were randomly divided into the experimental group (NKB group) and the control group (Saline group) and intravenously injected with NKB cells (1×10^5^ cells/mouse) or saline on day 5 and day 10 twice. (**B**) The arthritis scores in each group were recorded daily after the second immunization (n = 10 per group, two-way repeated-measures ANOVA test, \*\**P* < 0.01). (**C-H**) 19 days after the second immunization, the mice were sacrificed for further analyses. (**C**) Micro-CT images showing the bone destruction of paws from the mice of the CIA-NKB group, CIA- saline group, and naïve group. Arrows indicate bone destruction. (**D**) H&E staining of the sagittal sections of paws from each group of mice. Arrows indicate the stenosis the articular cavity and destruction of cartilage. Scale bar = 100 μm. (**E-H**) Th17 cells (**E, F**) and Th1 cells (**G, H**) were detected in the spleen of CIA-NKB, CIA-saline, and naïve mice by flow cytometry. The representative flow charts (**E, G**) and the statistical results (**F, H**) were shown, respectively (one-way ANOVA test followed by Tukey’s posttest for multiple comparisons, \**P* < 0.05, \*\**P* < 0.01, NS, not significant). Data are presented as mean ± SEM. Results are representative of three independent experiments.

**Figure 4.**
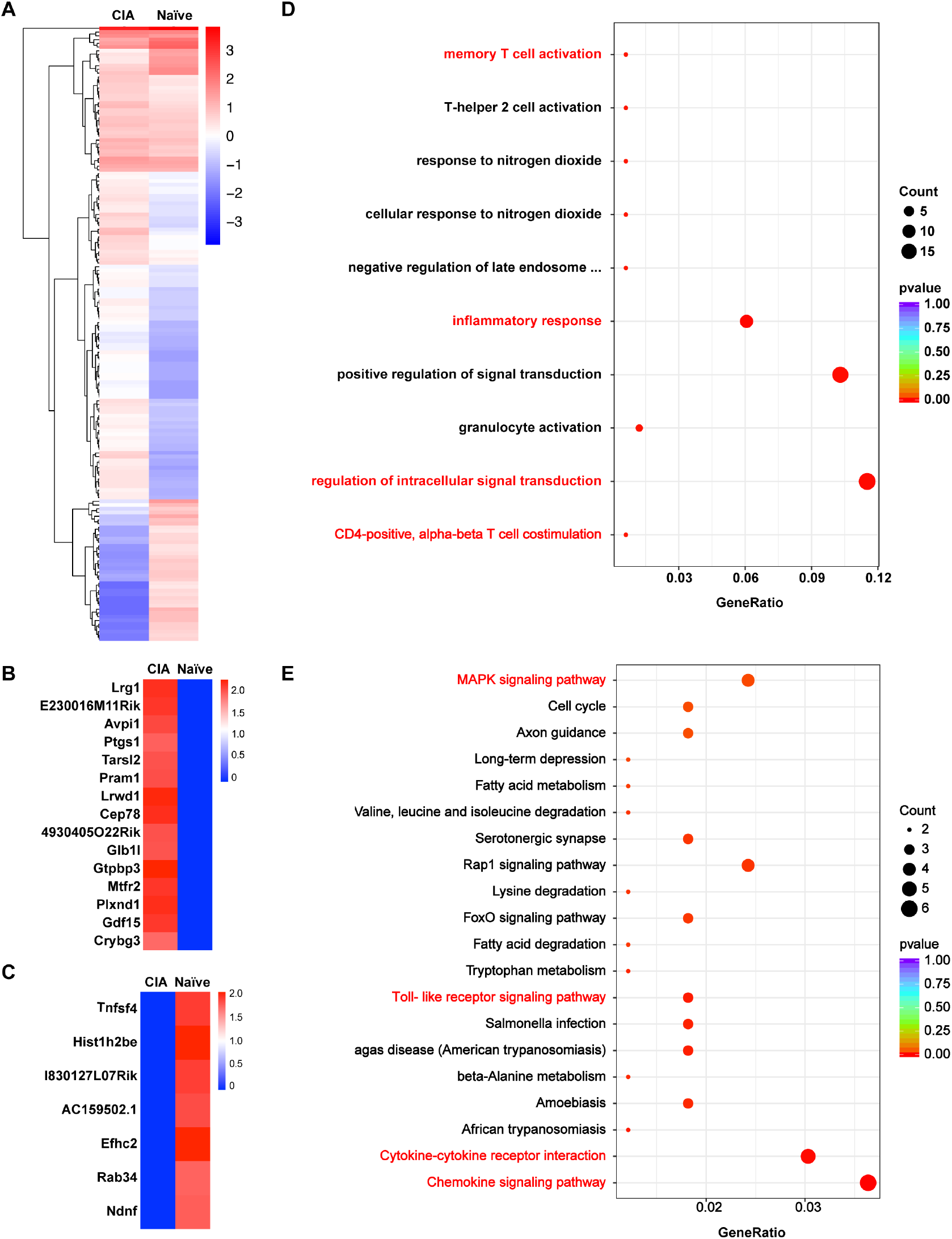
RNA sequencing of NKB cells from CIA and naïve mice. (**A**) Clustering analysis of the differentially expressed (DE) genes of NKB cells in CIA and naïve mice (n = 3 per group). Red and blue represent high and low expression levels, respectively. (**B**) Heat map of gene clustering specifically expressed in NKB cells of CIA mice. (**C**) Heat map of gene clustering specifically expressed in NKB cells of naïve mice. (**D**) Gene ontology (GO) analysis of DE genes. The biological process (BP) was shown. (**E**) KEGG pathway analyses of DE genes. Those ontologies and pathways associated with RA development were shown in red.

### NKB cells demonstrate differential gene expression profile in RA

To reveal the underlying mechanisms of NKB cells in aggravating RA development, we then illustrated the gene signature alteration of NKB cells under RA circumstance by RNA sequencing. 165 differentially expressed (DE) transcripts were identified, with 116 upregulated and 49 downregulated in CIA mouse NKB cells (Padj < 0.05, at least two-fold changes). These genes clearly distinguish CIA mice from naïve mice which were visualized in a hierarchical clustering diagram (Figure 4A). In particular, 15 genes were uniquely expressed in CIA mouse NKB cells (Figure 4B), while 7 genes were uniquely expressed in naïve mouse NKB cells (Figure 4C).

To explore the potential functional differences and molecular pathways of NKB cells in CIA and naïve mice, we categorized the DE genes according to the three GO systems: molecular function, biological process, and cellular component. These DE genes were found to be enriched in ontologies of CD4^+^ αβ T cell costimulation, memory T cell activation, regulation of intracellular signal transduction, and inflammatory response that potently promote RA progression (Figure 4D). KEGG pathway analysis further revealed the enrichment of these DE genes in the chemokine signaling pathway, cytokine-cytokine receptor interaction pathway, Toll-like receptor signaling pathway, and MAPK signaling pathway involved in RA pathogenesis (Figure 4E). It should be noted that IL-17C that promotes Th17 cell differentiation was upregulated while IL-10 that inhibits Th17 cell function(Chang et al., 2011; M. Yang et al., 2012) was downregulated in NKB cells under the arthritis condition (Supplementary Figure 5). Nevertheless, these key DE genes and their role in RA need to be further validated and investigated.

### Expansion of NKB cells in RA patients correlates with disease manifestation

Given the pathogenic role of NKB cells in CIA mice, we further detected their frequencies and analyzed the clinical associations in RA patients. The gating strategy for human NKB cells was displayed in Figure 5A. The results showed that compared with osteoarthritis (OA) patients and healthy controls, RA patients demonstrated significantly increased frequencies of NKB cells in the peripheral blood (Figure 5B, C). The more apparent expansion of NKB cells was also observed in RA patient synovial fluid (Figure 5D, E). Correlation analysis showed that the frequencies of NKB cells were positively correlated with RA patient RF and IgM levels, as well as the serum IL- 17A level, but not the ESR, CRP, anti-CCP, tender joint count (TJC), swollen joint count (SJC), and DAS28 (Figure 5F-I, Table 2). Moreover, after effective therapy with disease remission, RA patients revealed fundamentally declined frequencies of NKB cells to almost normal levels (Figure 5J-L). All these results suggest that the expansion of NKB cells in RA patients is actively involved in the disease progression.

**Figure 5.**
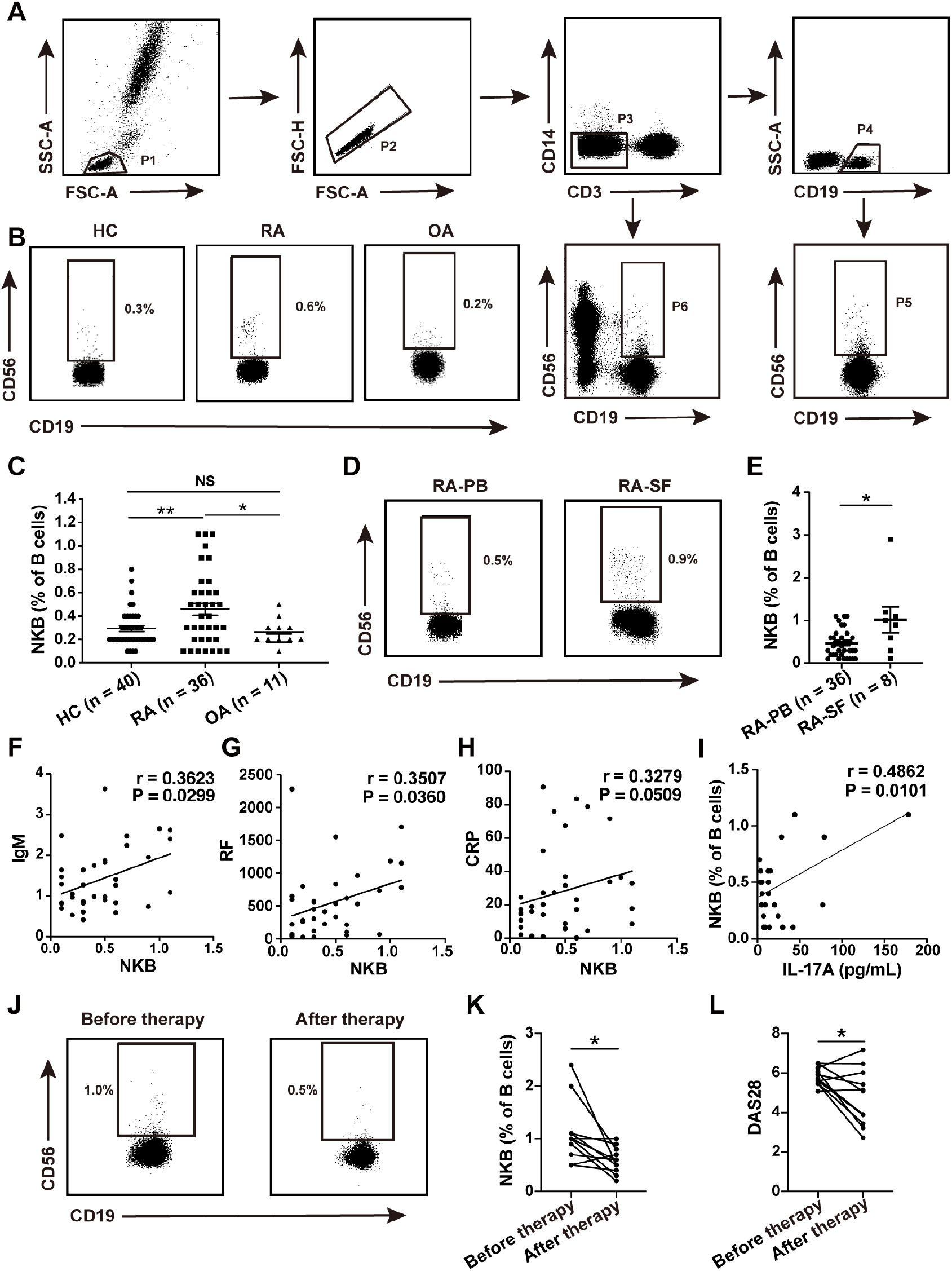
Expansion of NKB cells in RA patients. (**A**) Gating strategy for human NKB cells. (**B-C**) Flow cytometric analysis of the frequencies of NKB cells in the peripheral blood of 36 RA patients, 11 OA patients, and 40 healthy controls (HC). The representative flow charts (**B**) and the statistical result (**C**) were shown, respectively (one-way ANOVA followed by Tukey’s posttest for multiple comparisons, \**P* < 0.05, \*\**P* < 0.01, NS, not significant). (**D-E**) Flow cytometric analysis of the frequencies of NKB cells in the synovial fluid (n = 8) and peripheral blood (n = 36) of RA patients. The representative flow charts (**D**) and the statistical result (**E**) were shown, respectively (two-tailed Mann–Whitney *U* test, \**P* < 0.05). (**F-I**) The correlation of NKB cells with RA patient IgM (**F**), RF (**G**), CRP (**H**), and serum IL-17A(**I**) was analyzed, respectively (Spearman’s rank correlation test or Pearson correlation test). (**J-L**) 11 RA patients before (0W) and after therapy (12W) with disease-modifying antirheumatic drugs (DMARDs) were enrolled in the study. **(J, K**) The frequencies of NKB cells were detected by flow cytometry. The representative flow charts (**J**) and statistical result (**K**) were shown, respectively (two-tailed Wilcoxon signed-rank test, \**P* < 0.05). (**L**) The disease activity scores 28 (DAS28) of RA patients before and after therapy were evaluated and analyzed (two-tailed Wilcoxon signed-rank test, \**P* < 0.05). Data are presented as mean ± SEM.

**Table 1.** Clinical characteristics of RA patients.

| Characteristics | RA (n = 36) |
| --- | --- |
| Age, mean (range), yrs | 60.61 (29-80) |
| Sex, no. male/female | 5/31 |
| Duration, mean (range), yrs | 16.06 (1-42) |
| ESR, mean (range), mm/hr | 52.89 (5-107) |
| CRP, median (range), mg/l | 18.33 (0.22-90.54) |
| IgA, mean (range), g/l | 3.19 (1.02-6.67) |
| IgG, mean (range), g/l | 13.55 (5.4-22.3) |
| IgM, median (range), g/l | 1.18 (0.42-2.65) |
| RF, median (range), IU/ml | 429.5 (24.2-2280) |
| Anti-CCP, median (range), U/ml | 201.75 (49.69-257.03) |
| WBC, median (range), 10<sup>9</sup>/l | 5.6 (3.4-11.0) |
| HGB, mean (range), g/l | 111.18 (77-133) |
| PLT, mean (range), 10<sup>9</sup>/L | 249.68 (130-428) |
| TJC, median (range) of 28 joints | 5 (0-28) |
| SJC, median (range) of 28 joints | 3 (0-26) |
| DAS28, mean (range) | 5.10 (2.17-7.81) |
| Albumin, mean (range), g/l | 34.63 (27.6-41.4) |
| Medication, no. (%) |  |
| Steroids | 18 (50%) |
| NSAIDs | 19 (52.78%) |
| DMARDs | 34 (94.44%) |
| Biologics | 6 (16.67%) |
| No treatment | 0 (0%) |

ESR, erythrocyte sedimentation rate; CRP, C-reactive protein; IgA, immunoglobulin A; IgG, immunoglobulin G; IgM, immunoglobulin M; RF: rheumatoid factor; Anti- CCP: anti-cyclic citrullinated peptide antibody; WBC, white blood cell; HGB, hemoglobin; PLT, platelet; TJC, tender joint count; SJC, swollen joint count; DAS28: disease activity score 28; NSAIDs, nonsteroidal anti-inflammatory drugs; DMARDs, disease-modifying anti-rheumatic drugs.

**Table 2.**
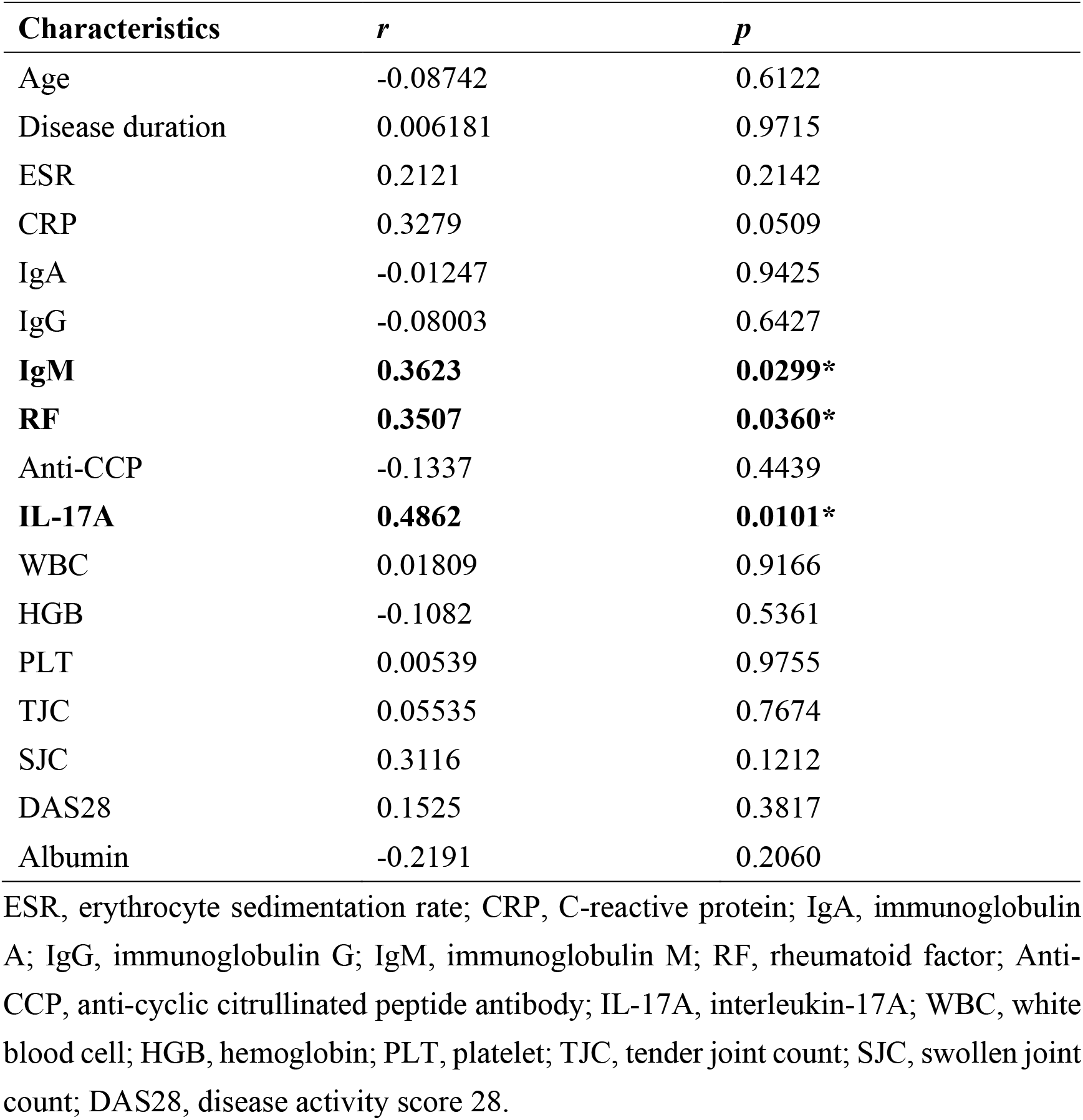
Association between NKB cells and RA patient clinical/immunological features.

## Discussion

In this study, we found that NKB cells were expanded both in arthritis mice and RA patients, positively correlated with the disease features. These cells could promote CD4^+^ effector T cell proliferation and Th17 cell differentiation, and exacerbate arthritis severity. Mechanically, NKB cells displayed distinct differential gene expression profiles under RA circumstances, potentially perpetuating the disease progression. Moreover, after effective therapy, NKB cells could be recovered to normal levels.

NKB cells were identified in the spleen and MLNs of both mice and human in the original study(Wang et al., 2016). The existence of NKB cells was questioned by some researchers after it was published(Kerdiles et al., 2017), but Shuo Wang and the co- authors came up with a well-founded response that confirmed NKB cells as a separate subset of innate B cells(Wang et al., 2017). Moreover, the following study showed that NKB cells were widely distributed in rhesus monkeys, including spleen, peripheral blood, tonsil, lymph node, colon, and jejunum. Besides, they also found that NKB cells existed in the peripheral blood of human(Manickam et al., 2018). In our study, we found that NKB cells were systematically distributed in DBA/1 mice, including spleen, MLNs, DLNs, Peyer patch, and paw, with spleen as the dominant enriching site. In addition, we also identified NKB cells in the peripheral blood and synovial fluid of healthy individuals and RA patients. Due to the limitation of sample sources, we only detected NKB cells in human peripheral blood and synovial fluid, but the frequencies of NKB cells in tissues (such as spleen and lymph nodes) should be higher.

NKB cells were proved to prime innate lymphocytes against microbial infection, and participate in the SIV infection and experimental alcoholic liver injury. Nevertheless, their role in autoimmune diseases remains completely unknown. In the present study, we found that NKB cells were significantly expanded both in arthritis mice and RA patients. Moreover, these cells could perpetuate the inflammation and exacerbate the disease severity of RA. Our data preliminarily revealed the pathogenic role of NKB cells in RA, which needs to be further studied in other autoimmune diseases.

The mechanisms of NKB cells in the disease pathogenesis remain largely unclear. Here, we showed that NKB cells could promote CD4^+^ effector T cell proliferation in RA. Moreover, NKB cells potently primed Th17 cell differentiation, one of the key effector cells in RA pathogenesis(Azizi, Jadidi-Niaragh, & Mirshafiey, 2013; Hirota et al., 2018; J. Yang, Sundrud, Skepner, & Yamagata, 2014; P. Yang et al., 2019), thus exacerbating the disease progression. Both cell-cell contact and soluble mediators were proved to be involved in NKB cell-mediated Th17 cell proinflammatory responses. Nevertheless, the key soluble mediators were undetermined. Previous study showed that NKB cells expressed several cytokines, including IL-1β, IL-6, IL-12, IL-15, and IL-18. IL-18 was found to be a signature cytokine of NKB cells and played an indispensable role in NKB cells-mediated eradication of microbial infection(Wang et al., 2016). IL-6 is well recognized in Th17 cell differentiation(Harbour et al., 2020). Whether IL-18 or other cytokines derived from NKB cells were also involved in the enhanced Th17 cell responses needs to be further investigated. In addition, we also found that NKB cells were positively correlated with RA patient IgM, RF, as well as serum IL-17A level. This indicated that NKB cells might be involved in the pathogenesis of RA by participating in Th17 cell proinflammatory responses and producing autoantibodies, which need to be further confirmed. Due to the low frequencies of NKB cells and the limited volume of blood samples denoted by the healthy volunteers and RA patients, we failed to obtain enough NKB cells for functional studies. Besides, whether NKB cells participate in osteoclast formation and osteoblast inhibition merits in-depth studies.

In summary, here we revealed the expansion and pathogenic role of NKB cells in RA by promoting Th17 cell differentiation. These cells might be served as a potential therapeutic target for RA.

## Materials and methods

### Patients and controls

55 RA patients, 11 OA patients, and 40 healthy controls were enrolled in the study. All RA patients fulfilled the 2010 American College of Rheumatology (ACR) and European League Against Rheumatism (EULAR) criteria for RA. All OA patients fulfilled ACR 1995 criteria for OA. The study was approved by the Institutional Medical Ethics Review Board of Peking University People’s Hospital, and all the participants provided written informed consent.

### Mice

Six to eight-week-old male DBA/1 mice were obtained from Huafukang Bioscience Company (Beijing, China). All mice were housed in a specific pathogen-free environment and were acclimatized for a week before the experiment. All animal procedures were approved by the institutional animal care and use committee (IACUC) of Peking University People’s Hospital.

### Experimental arthritis model

CIA models were established in DBA/1 mice by immunization with bovine type II collagen (CII, Chondrex, Redmond, WA). Briefly, DBA/1 mice were first immunized with 200 μg CII emulsified with Freund’s complete adjuvant (CFA, Chondrex) on day 0, and were secondly immunized with 100 μg CII emulsified with Freund’s incomplete adjuvant (IFA, Chondrex) on day 21. The severity of arthritis was scored based on the inflammation level in each of the four paws as previous description(F. Hu et al., 2020).

### Isolation of NKB cells

CD49b^+^ cells from splenocytes of CIA mice were first enriched by Magnetic Cell Sorting system (Mouse CD49b Positive Selection Kit, STEMCELL, Canada), then were stained with FITC-anti-mouse CD19 antibody (Biolegend, San Diego, CA) and sorted on FACS Arial II flow cytometer (Becton Dickinson, San Diego, CA). The purity of the isolated NKB cells (CD19^+^CD49b^+^) was over 90% as detected by flow cytometry. The isolated NKB cells were then subjected to *in vitro* and *in vivo* functional studies.

### CD4^+^ effector T cell proliferation assay

The proliferation of CD4^+^CD25^-^ effector T cells was measured by CFSE (CellTrace™ CFSE Cell Proliferation Kit, Thermal Fisher Scientific, Waltham, MA) dilution assay. Briefly, 5×10^4^ CFSE labeled effector T cells and different numbers of NKB cells (1×10^4^, 5×10^4^) from CIA mice were co-cultured, respectively, in the presence of plate- bound anti-CD3 (5 μg/mL, Biolegend), CpG (10 μg/mL, Sangon Biotech, Beijing, China), anti-CD28 and anti-CD40 (3 μg/mL, Biolegend) in a round-bottom 96-well plate. Three days later, the cells were collected and stained with APC-Fire750-anti- mouse CD4 antibody (Biolegend), then were detected by flow cytometry. The frequencies of proliferating cells were determined by analyzing CFSE^low/-^ cells in CD4^+^ T cells.

### CD4^+^CD25^-^ effector T cells: NKB cells co-culture assay

5×10^5^ CD4^+^CD25^-^ T cells were co-cultured with 1×10^5^ NKB cells (5:1, both from CIA mice) under normal or transwell culture systems in the presence of plate-bound anti- CD3, CpG, anti-CD28 and anti-CD40 as previous describe. Three days later, the cells were harvested for flow cytometry, while the cell culture supernatants were collected for ELISA.

### Th17 cell differentiation assay

5×10^5^ CD4^+^CD25^-^CD62L^hi^ naïve T cells and 1×10^5^ NKB cells from CIA mice were co-cultured in the presence of plate-bound anti-CD3 (5 μg/mL), anti-CD28 (1 μg/mL), TGF-β (2.5 ng/mL, Peprotech, Rocky Hill, NJ), IL-6 (20 ng/mL, Peprotech), anti-CD40 (3 μg/mL), CpG (10 μg/mL), anti-IL-4 and anti-IFN-γ (5 μg/mL, Biolegend) for 3 days.

Then, the cells were harvested for Th17 cell detection, while the cell culture supernatants were collected for IL-17A analysis.

### NKB cells adoptive transfer assay

The onset stage CIA mice with an average arthritis score of 2 were randomly divided into the experimental group (NKB group) and the control group (Saline group). Then, the mice were intravenously injected with NKB cells from CIA mice (1×10^5^ cells/mouse) or saline on day 5 and day 10 post the second immunization twice. The arthritis scores in each group were recorded daily. 19 days after the second immunization, the mice were sacrificed for flow cytometry analysis, micro CT scan, and H&E staining.

### Flow cytometry analysis

For surface staining, cells were stained with various antibodies at room temperature for 30 min, washed with PBS, and then re-suspended and fixed with 2% formaldehyde under room temperature. For intracellular staining, cells were incubated with PMA, ionomycin, and BFA for 5 h, then surface stained, fixed, permeabilized, and intracellular stained according to the manufacture’s instruction. Cells were analyzed on FACS Arial II and CytoFLEX (Beckman Coulter, Brea, CA) flow cytometers.

### ELISA analysis of IL-17A and IFN-γ

The cell culture supernatants or serum were collected and stored at -80°C until use. Commercial available mouse/human IL-17A and IFN-γ ELISA kits (NEOBIOSCIENCE, Beijing, China) were used to detect the cytokines according to the manufacturer’s instructions. The results were measured on the Synergy^TM^ 4 Multi- Mode Microplate Reader with software GEN5CH 2.0 (BioTek, Winooski, VT).

### Mouse paw histopathology and micro-CT

For histological analysis and micro-CT scan, mice were sacrificed and paws were collected and fixed in 4% buffered formaldehyde. Paws were then paraffin-embedded, sectioned, stained with H&E, and analyzed with NDP.view2 (Hamamatsu Photonics K.K., Japan). Micro-CT images were acquired on the Tri-Modality FLEX Triumph™ Pre-Clinical Imaging System (Gamma Medica-Ideas, Northridge, CA). CT image sets acquisitions lasted 10 min and utilized beam parameters of 130 μA and 80 kVP. Analyze 10.0 (AnalyzeDirect, Overland Park, KS) was used to perform the image analysis.

### RNA-seq

Isolated NKB cells from 3 active CIA mice and 3 naïve mice (∼1,000 cells per sample) were subjected to RNA sequencing accomplished by Beijing Annoroad Gene Technology Co., Ltd. (Beijing, China). The significantly differentially expressed (DE) genes (Padj < 0.05, at least two-fold changes) were first set to hierarchical cluster analysis. To reveal their biological significance, these DE transcripts were further categorized according to the gene ontology (GO) and Kyoto Encyclopedia of Genes and Genomes (KEGG) databases.

### Statistics

All statistical calculations were performed using the statistical software programs SPSS 26.0 (SPSS Statistics Inc.) and Graphpad prism 8 (GraphPad Software Inc., San Diego, CA). Differences between various groups were evaluated by the student’s *t*-test, one-way ANOVA test, Mann–Whitney *U* test, paired *t*-test, Wilcoxon signed-rank test, Spearman’s rank correlation test, or two-way repeated-measures ANOVA test, and were considered statistically significant at *P* < 0.05.

## Acknowledgments

This work was supported by grants from the National Natural Science Foundation of China (81971523, 81671604, and 81302554 to Dr. F. Hu, U1903210 to Dr. Z. Li, 81871281 to Dr. Y. Jia, and 81871289 to Dr. H. Ye), the Beijing Nova Program (Z181100006218044 to Dr. F. Hu), and Beijing Science and Technology Planning Project (Z191100006619111 to Dr. Y. Su, and Z191100006619109 to Dr. H. Ye), as well as by the Fundamental Research Funds for the Central Universities: Peking University Clinical Medicine Plus X-Young Scholars Project (PKU2021LCXQ014 to Dr. F. Hu), and the Peking University People’s Hospital Research and Development Funds (RDX2020-01 to Dr. F. Hu). The funders had no role in study design, data collection, and analysis, decision to publish, or preparation of the manuscript.

## Contributions

Performed the experiments: P.W., J.S., and M.X.B.; Analyzed the data: P.W., X.Z., Y.X., Y.D.T., X.L., and X.Y.F.; Contributed reagents/materials/analysis tools: Y.J., L.M.R., H.Y., Y.S., S.W.; Wrote the manuscript: P.W.; Reviewed and edited the manuscript: Z.G.L. and F.L.H.

## Supplementary material

**Figure S1.**
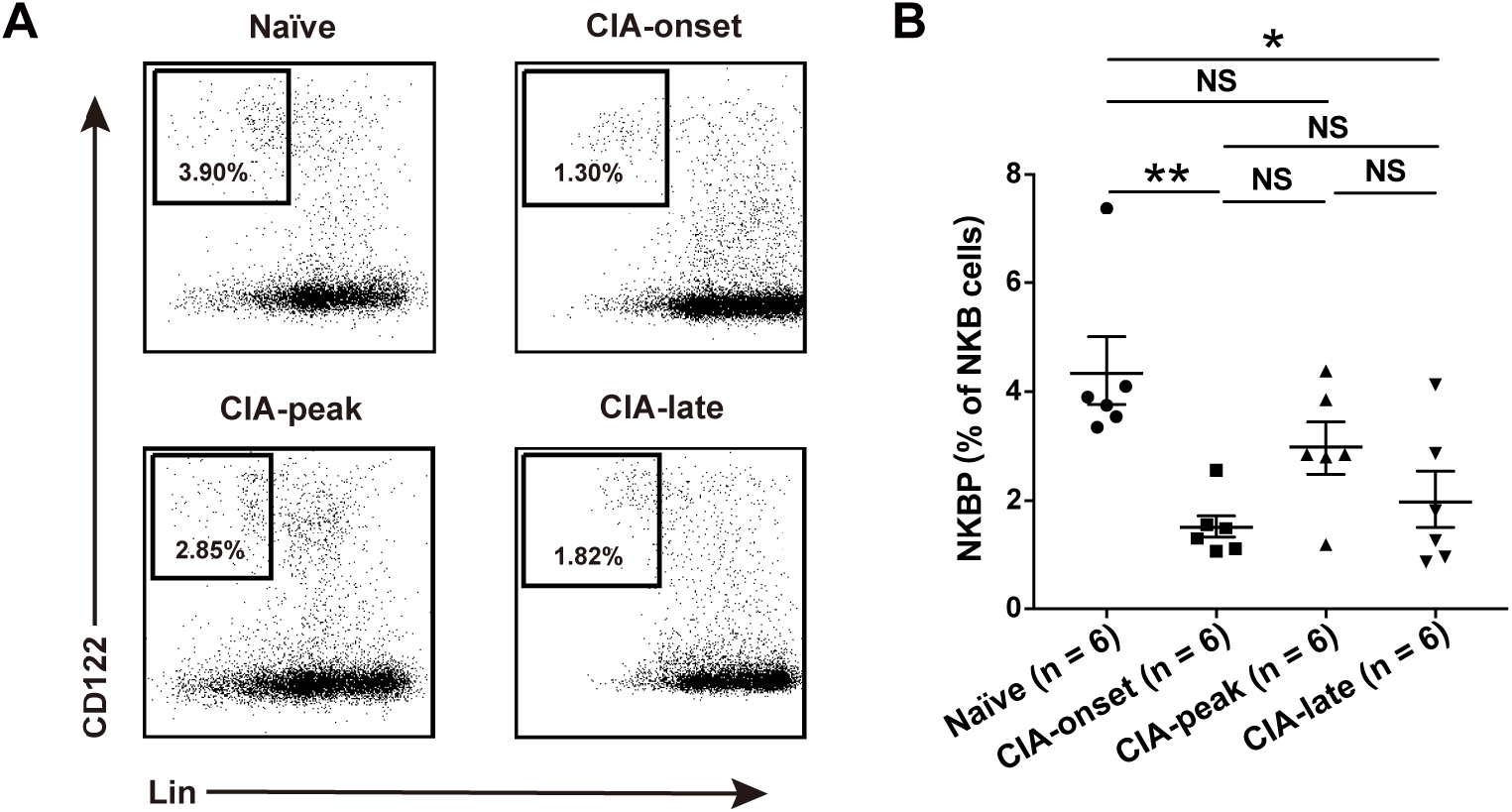
The dynamic changes of NKB cell precursors (NKBP) during the development of collagen-induced arthritis (CIA). Flow cytometric analysis of the frequencies of NKBP during the development of CIA. Spleens from naïve, CIA-onset (< 2 days), CIA-peak (∼ 15 days), and CIA-late (> 60 days) were collected for NKBP detection (n = 6 per group). The representative flow charts (**A**) and the statistical result (**B**) were shown, respectively (one-way ANOVA test followed by Tukey’s posttest for multiple comparisons, \**P* < 0.05, \*\**P* < 0.01, NS, not significant). Data are presented as mean ± SEM. Results are representative of three independent experiments.

**Figure S2.**
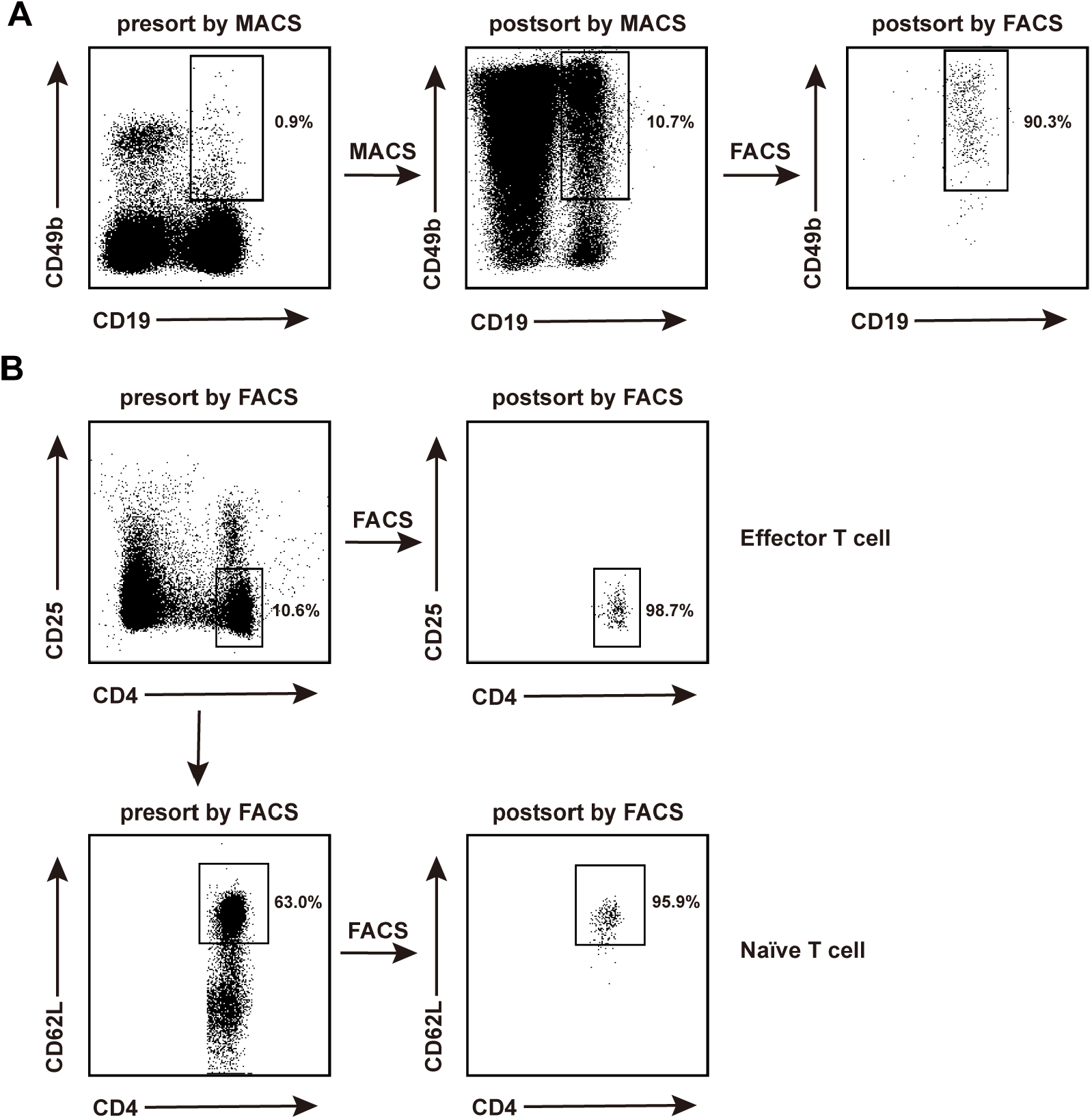
The representative flow charts of purity of NKB cells, effector T cells, and naïve T cells. (**A**) CD49b^+^ cells from splenocytes of CIA mice were first enriched by Magnetic Cell Sorting system (MACS), then were stained with FITC-anti-mouse CD19 antibody and sorted on FACS Arial II flow cytometer. The purity of the isolated NKB cells (CD19^+^CD49b^+^) was detected by flow cytometry. (**B**) CD4^+^CD25^-^ effector T cells and CD4^+^CD25^-^CD62L^hi^ naïve T cells from splenocytes of CIA mice were respectively sorted on FACS Arial II flow cytometer, and the purities of which were detected by flow cytometry.

**Figure S3.**
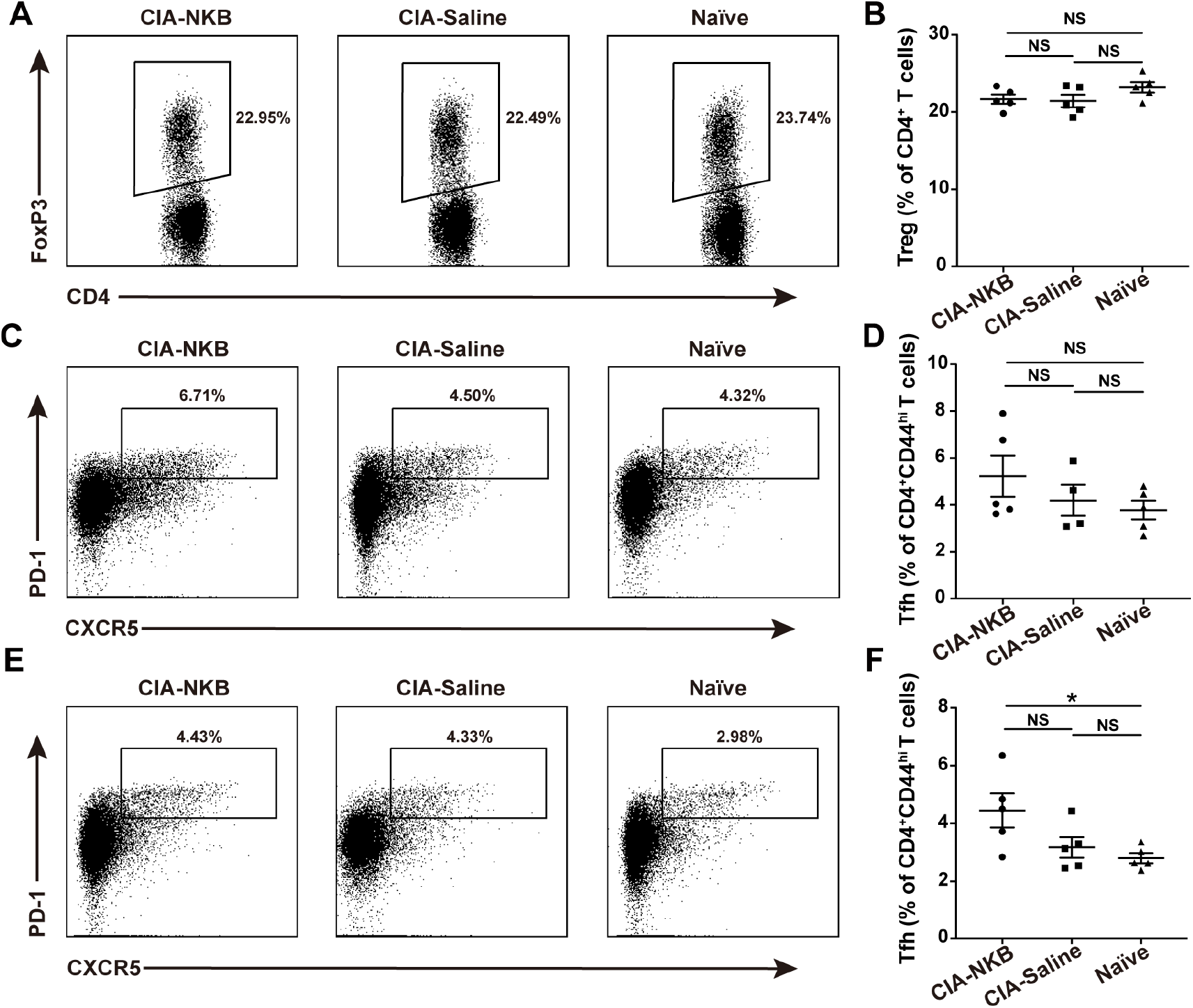
Alteration of Treg and Tfh cells in CIA mice after adoptive transfer of NKB cells. Flow cytometric analysis of the frequencies of Treg (spleen: **A, B**) and Tfh (DLN: **C, D**; MLN: **E, F**) cells in CIA mice after adoptive transfer of NKB cells or saline. The representative flow charts (**A, C, E**) and the statistical results (**B, D, F**) were shown, respectively (one-way ANOVA test followed by Tukey’s posttest for multiple comparisons, \**P* < 0.05, NS, not significant). Data are presented as mean ± SEM. Results are representative of three independent experiments.

**Figure S4.**
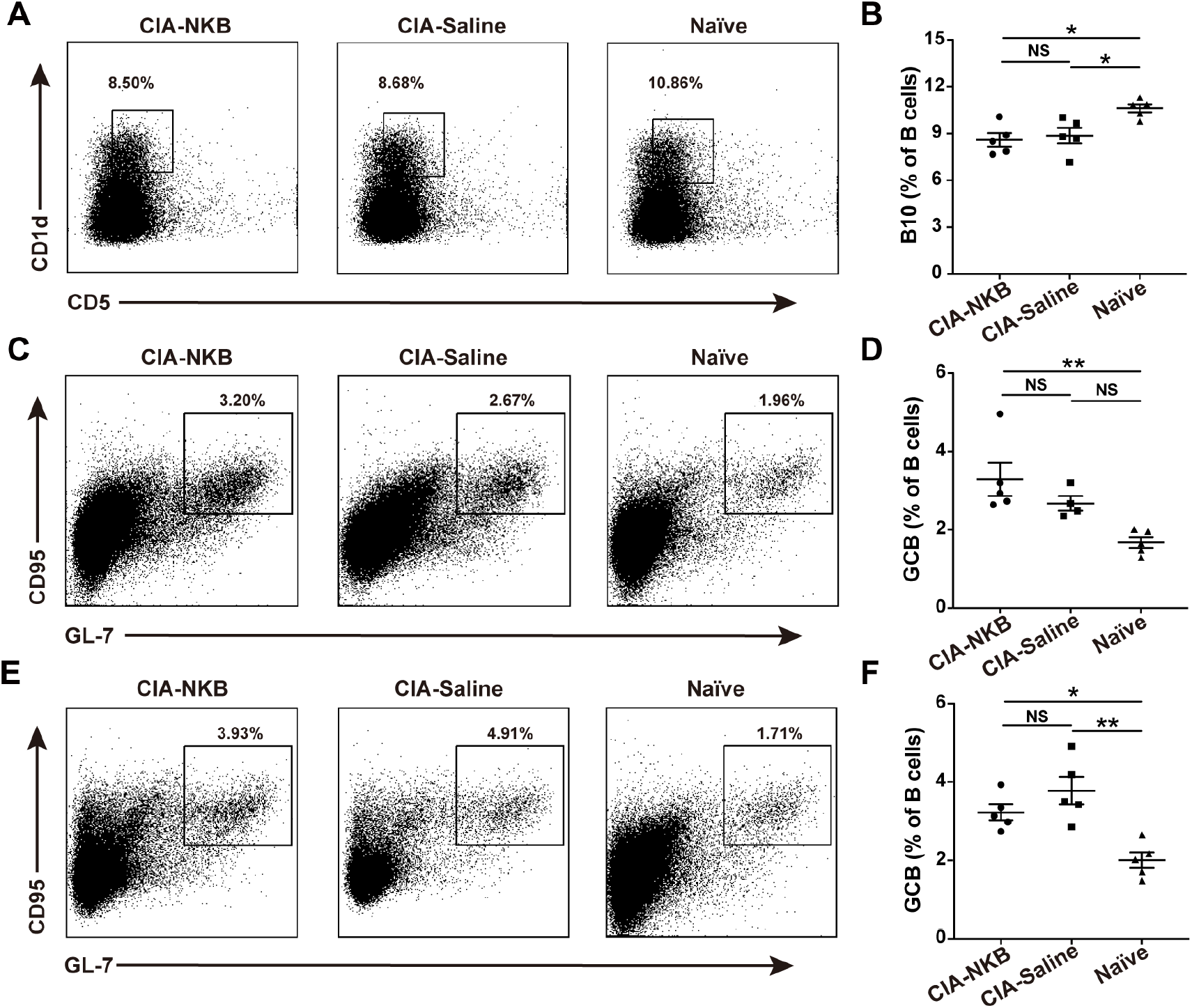
Alteration of B10 and Germinal center B (GCB) cells in CIA mice after adoptive transfer of NKB cells. Flow cytometric analysis of the frequencies of B10 cells (spleen: **A, B**) and GCB (DLN: **C, D**; MLN: **E, F**) cells in CIA mice after adoptive transfer of NKB cells or saline. The representative flow charts (**A, C, E**) and the statistical results (**B, D, F**) were shown, respectively (one-way ANOVA test followed by Tukey’s posttest for multiple comparisons, \**P* < 0.05, \*\**P* < 0.01, NS, not significant). Data are presented as mean ± SEM. Results are representative of three independent experiments.

**Figure S5.**
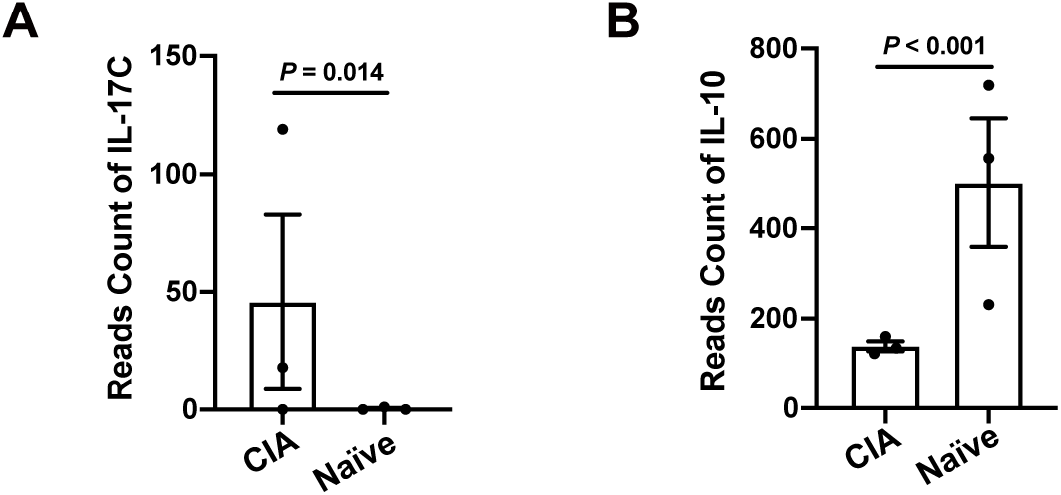
Different gene expression of IL-17C and IL-10 in NKB cells between CIA mice and naïve mice. (A, B) The statistical results of gene expression of IL-17C **(A)**, and IL-10 were shown **(B)**, respectively (Linear regression was used to estimate the expression intensity of genes in each sample, and Wald test was used to calculate the p-value of probability of non-differential expression of each gene in the two groups of samples). Data are presented as mean ± SEM.

